# Selective binding and transport of protocadherin 15 isoforms by stereocilia unconventional myosins in a heterologous expression system

**DOI:** 10.1101/2022.02.25.482032

**Authors:** Angela Ballesteros, Manoj Yadav, Runjia Cui, Kiyoto Kurima, Bechara Kachar

**Author notes:** These authors contributed equally (Angela Ballesteros, Manoj Yadav). Corresponding authors: Angela Ballesteros, Ph.D., Current address: Molecular Physiology and Biophysics Section, NINDS, NIH, Bethesda, MD 20892. and Bechara Kachar, M.D., Laboratory of Cell Structure and Dynamics, NIDCD, NIH, Bethesda, MD 20892, USA.

## Abstract

During hair cell development, the mechanoelectrical transduction (MET) apparatus is assembled at the stereocilia tips, where it coexists with the stereocilia actin regulatory machinery. While the myosin-based tipward transport of actin regulatory proteins is well studied, isoform complexity and built-in redundancies in the MET apparatus have limited our understanding of how MET components are transported. We used a heterologous expression system to elucidate the myosin selective transport of isoforms of protocadherin 15 (PCDH15), the protein that mechanically gates the MET apparatus. We show that MYO7A selectively transports the CD3 isoform while MYO3A and MYO3B transports the CD2 isoform. Furthermore, MYO15A showed an insignificant role in the transport of PCDH15, and none of the myosins tested transport PCDH15-CD1. Our data suggest an important role for MYO3A, MYO3B, and MYO7A in the MET apparatus formation and highlight the intricate nature of MET and actin regulation during development and functional maturation of the stereocilia bundle.

## INTRODUCTION

The hallmark of hair cells, the sensory receptors of the inner ear is their mechanosensitive organelle, the hair cell bundle. This bundle consists of rows of actin-based protrusions of graded heights called stereocilia forming a characteristic staircase architecture. The hair cell bundle converts mechanical stimuli into electrical signals through a mechanoelectrical transduction (MET) apparatus, built around a protein filament, the tip-link, that connects the tip of the shorter stereocilia to its closest taller neighbor^1^. This sophisticated architecture ensures the optimal gating of the MET channels located at the lower end of the tip-link, when the stereocilia are deflected toward the taller stereocilia^2^. Remarkably, the MET molecular apparatus co-exists at stereocilia tips with the actin regulatory protein complex that sustains the stereocilia length differential, and there is increasing evidence of coordination between these two molecular machineries^3–5^.

Stereocilia tips can be up to a staggering 100 μm away from the hair cell body, which together with the restricted space between the stereocilia actin core and the plasma membrane makes it unlikely that molecular components efficiently traverse from cell body to stereocilia tips by simple diffusion. Like in other actin protrusions, active transport of molecular components in stereocilia is carried out by unconventional myosins, monomeric or dimeric myosins that transport cargoes along the surface of actin filament bundles^6^. The unconventional myosins MYO3A, MYO3B, and MYO15A have been implicated in transporting different actin-regulatory and scaffolding protein cargo to stereocilia tips^7–9^, but little is known about the transport of the MET components to stereocilia tips. Mutations in the MYO3A, MYO3B, MYO7A, and MYO15A genes have been implicated in different forms of inherited hearing loss through specific disruption of stereocilia structure and its normal mechanosensitivity^10–17^, suggesting a role for these myosins in stereocilia architecture and MET. The appearance of MYO3A in auditory stereocilia coincides with the onset of MET^18^, suggesting a role for MYO3 in the transport of the MET apparatus. Additionally, MYO7A localize to the stereocilia tips and has been implicated in regulating stereocilia structure and MET function^19–21^. Stereocilia myosins, like all myosins, typically share a similar motor domain but differ in their cargo-binding tail domain^22^. Because of the multitude of cargo and myosin tail domains, we hypothesize that stereocilia myosins selectively transport MET components to stereocilia tips but leave open the possibility for partial complementary functions and redundancy. Sorting out the pairing of motor and cargo is fundamental to understand the trafficking and selective targeting of stereocilia tip components, including the MET components that are essential for hearing and balance. How MET components are targeted to the tips of a specific stereocilia row in the staircase is key to understanding the mechanisms of MET complex assembly as well as an alluring cellular biology question. A key component of the MET complex is protocadherin 15 (PCDH15), which makes up the lower portion of the tip-link that mechanically gates the MET channel^23,24^. Hair cells express three isoforms of PCDH15 (PCDH15-CD1, -CD2, and -CD3) that share a common extracellular domain but differ in the C-terminal region of their cytoplasmic tails^25^. The expression levels, localization, and role of each isoform is not fully understood but several studies have shown partial redundancy within the three PCDH15 isoforms during development with the CD2 isoform being essential for proper MET function in mature auditory hair cells^26–28^.

Studying transport of molecular components to the MET site at stereocilia tips is challenging because of the intricate relationships between molecular motors, actin regulation, and mechanosensitivity^7,9,19,20,29,30^. Perturbation of virtually any stereocilia component can either be masked by redundancy in function or rapidly cascade into disruption of hair bundle structure and function. Here we investigate the ability of myosins 3A, 3B, 7A, 10, and 15A, the prevalent myosins found in stereocilia, to individually transport the three main PCDH15 isoforms along filopodial actin protrusions in model cultured cells. We show that MYO3A and MYO7A show selective binding and transport of two distinct PCDH15 isoforms along actin protrusions. We argue that the selective transport of PCDH15 isoforms by stereocilia myosins is part of the dynamic mechanisms of spatiotemporal positioning of these key inter-stereociliary links and MET components to their functional location in the stereocilia.

## RESULTS

### Dynamic localization of stereocilia myosins and PCDH15 during hair cell bundle development

During development of the stereocilia bundle of auditory hair cells, PCDH15 isoforms show a tipward localization, and are presumed to be part of the stereocilia tip-links, radial links, and kinocilium-stereocilia links (Fig. 1a and Supplementary Fig. 1a). When the hair bundle matures, the radial links are pruned, the kinocilium is reabsorbed, and PCDH15 is mainly localized at the tips of the shorter stereocilia as part of the MET complex apparatus^26^. Vestibular hair cells follow a similar developmental transition, but their mature stereocilia bundles conserve the kinocilium (Fig. 1a and Supplementary Fig. 1b). The cargo-transporting stereocilia myosins 3A, 3B, 15A, and 7A also undergo changes in spatiotemporal localization during development and have been shown to dynamically localize at stereocilia tips (Fig. 1a) in mature bundles^7,18,20,31^. Of these myosins, MYO7A, presents a major localization change since it shows additional specialized accumulation near the stereocilia base and at the upper tip-link density in mature auditory hair cells^19,20,32^. To test for selective pairing of stereocilia myosins with PCDH15 isoforms as putative cargo, we examined their co-localization and dynamics along filopodia in cultured COS7 cells expressing combinations of these proteins.

**Figure 1:**
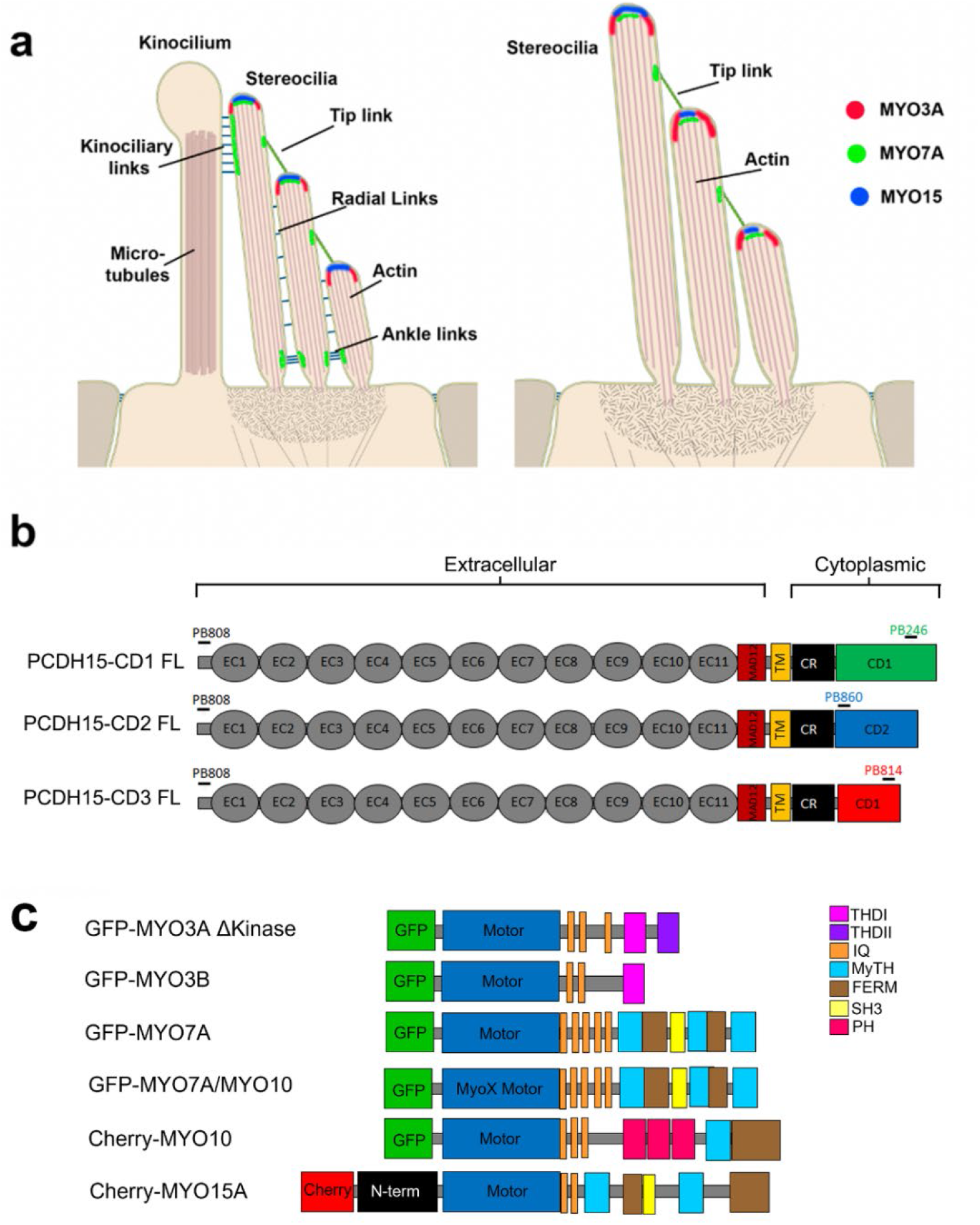
Localization of stereociliary myosin in the hair cell bundle and protein constructs used in this study. **a**) Localization of MYO3A (red), MYO7A (green), and MYO15 (blue) in the hair cell bundle during development or in the vestibular organs (left panel) and in mature auditory hair cells (right panel). **b**) Representation of the PCDH15 constructs used in this study. PCDH15 is a large cadherin protein with 11 extracellular calcium-binding (EC) domains, an extracellular linker or membrane adjacent domain (MAD12), a transmembrane ^65^ segment, and a cytoplasmic domain. The cytoplasmic domain contains a common region (CR) and a unique tail domain specific to each isoform (CD1, CD2, and CD3). Constructs containing the cytoplasmic (cyto) domain of PCDH15-CD1, -CD2 and -CD3 were also generated. **c**) Diagram of the myosin constructs used in this study.

### Generation and validation of PCDH15 constructs and antibodies

All three main PCDH15isoforms; PCDH15-CD1, -CD2, and -CD3, which differ in their cytoplasmic domain (Fig. 1b), are expressed in the hair cell bundle^25,27^. We obtained or cloned murine full-length (FL) and generated cytoplasmic (cyto) domain cDNA constructs for all three isoforms. To track the PCDH15 isoform localization, we generated affinity purified isoform-specific antibodies targeting peptide sequences unique to each PCDH15 isoform, as well as a common peptide sequence from the conserved ectodomain region (Table 1). We confirmed the affinity and specificity of these antibodies in COS7 cells expressing the untagged PCDH15 FL isoforms (Supplementary Fig. 2) or their respective cyto-domains fused with a GFP at their N-termini (Supplementary Fig. 3d). All four antibodies showed selective labeling of PCDH15-expressing cells, and we did not detect any cross-reactivity of the isoform-specific antibodies with the other isoforms (Supplementary Figs. 2 and 3).

**Table 1:** Custom made polyclonal anti PCDH15 antibodies used in this study. The name, specific PCDH15 isoform recognized by these antibodies and the protein peptide used to generate the rabbit anti PCDH15 antibodies is indicated.

| **Antibody** | **Protein epitope** | **Epitope sequence** |
| --- | --- | --- |
| PB808 | pan-PCDH15 | GQYDDDWQYEDCKLARGG |
| PB246 | PCDH15-CD1 | PASNPQWGAEPHRHPK |
| PB860 | PCDH15-CD2 | EGEKARKNIVLARRRP |
| PB814 | PDCD15-CD3 | AVKPSGTRLKHTAE |

The myosins used in this study were tagged at their N-termini with either GFP or mCherry (Fig. 1c). We used a MYO3A lacking its N-terminal regulatory kinase domain since it localizes more efficiently to filopodia tips^33,34^. Because native MYO7A full length ^35^ tends to fold and does not induce filopodia in COS7 cells, we used a chimera that has a myosin 10 (MYO10) motor and a MYO7A tail domain that has been shown to induce filopodia formation and transport cargo to filopodia tips in COS7 cells ^36^. MYO10 has been shown to be a rare stereocilia myosin ^37^, but we included it in our experiments as a control for the MYO10 (motor)-MYO7A (tail) chimera.

### MYO3A specifically and actively transports PCDH15-CD2 to the tips of filopodia

When MYO3A is expressed in COS7 cells it induces the formation of filopodia with MYO3A enriched at its tips, which provides a well-defined spatial compartment where potential interactions can be clearly visualized ^18^. Cells co-expressing MYO3A and cyto PCDH15 showed that PCDH15-CD2, but not CD1 or CD3, is efficiently targeted to the tips of filopodia initiated by MYO3A (Fig. 2a). We quantified the distribution of PCDH15 by calculating the filopodia tip to filopodia shaft fluorescence ratio as described in the Material and Methods section. We observed an increase in the filopodia tip to shaft ratio of PCDH15-CD2 while the expression of MYO3A did not produce any filopodia tip enrichment of either CD1 or CD3 isoforms (Fig. 2b). In addition, we quantified the enrichment of MYO3A at the filopodia tips by examining the filopodia tip to shaft fluorescence ratio. Interestingly, we observed that that the enrichment of MYO3A at the filopodia tips was significantly enhanced in the presence of the cyto PCDH15-CD2 but not in the presence of-CD1 and-CD3 isoforms (Fig. 2c), suggesting some form of cooperativity between the MYO3A motor activity and the PCDH15-CD2 cargo.

**Figure 2:**
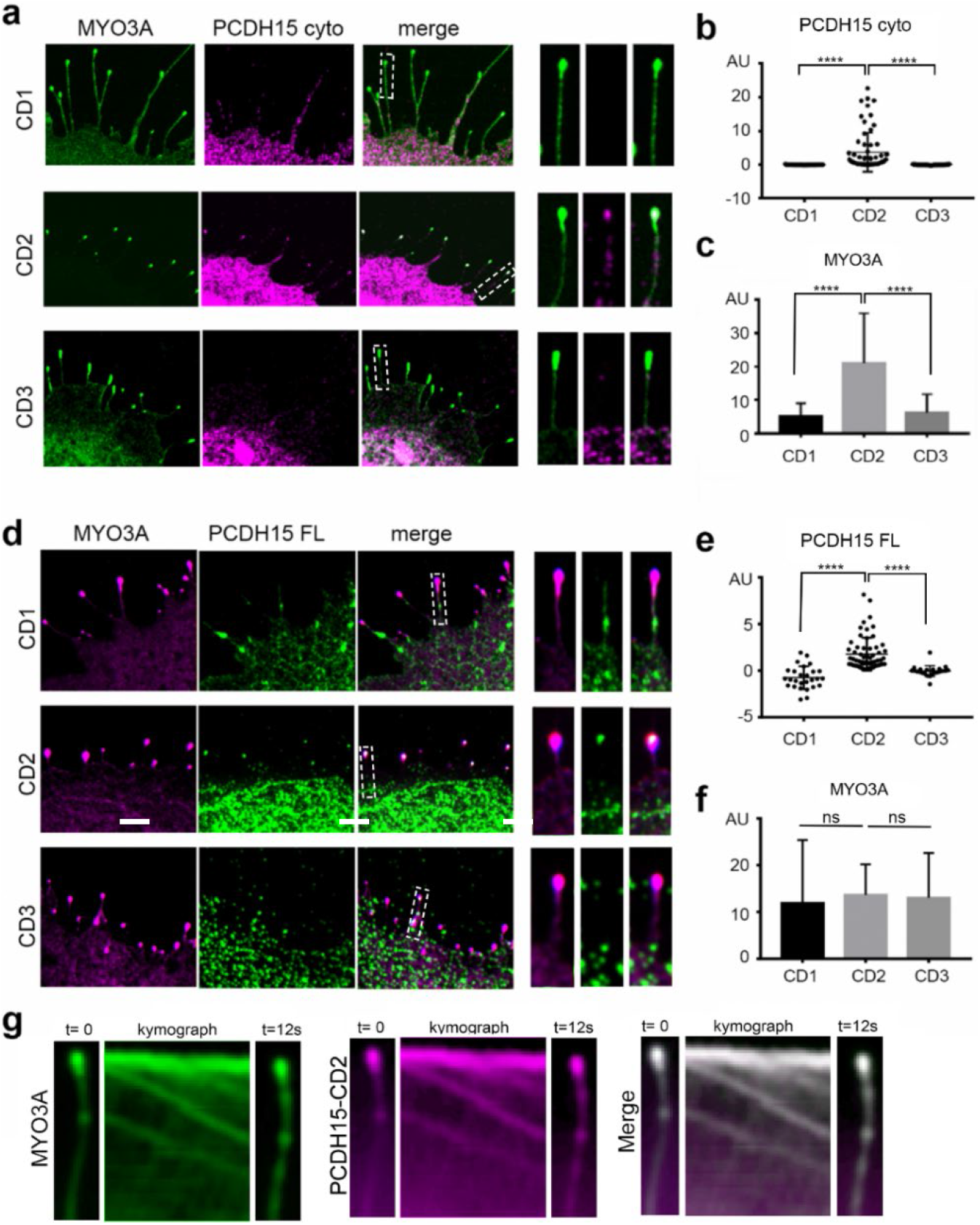
MYO3A actively transport PCDH15-CD2, but not CD1 or CD3, to the filopodia tips. **a**) Confocal images of COS7 cells expressing GFP-MYO3A △K (green) and the cytoplasmic domain (cyto) of PCDH15-CD1, -CD2, or -CD3 (magenta). The right panels show a close-up view of a representative filopodia. **b**) PCDH15-CD2 cyto exhibits significantly higher tip to cell ratio than PCDH15-CD1 or PCDH15-CD3 when co-expressed with MYO3A. **c)** Expression of PCDH15-CD2 enhanced the MYO3A enrichment at the filopodia tips (filopodia tip/shaft ratio). **d)** Confocal images of COS7 cells expressing GFP-MYO3A △K (magenta) and Full-length PCDH15-CD1, -CD2, or -CD3 (green). **e**) Full length PCDH15-CD2 exhibits significantly higher tip to cell ratio than PCDH15-CD1 or PCDH15-CD3 when co-expressed with MYO3A. **f**) MYO3A exhibit a similar enrichment at the filopodia tips when co-transfected with FL PCDH15-CD1, -CD2 or -CD3. Each dot in b and e represents one filopodia. One-way ANOVA analysis was performed with Dunnett’s multiple comparisons test. Level of significance was determined based on the p value (n.s p>0.05, *p<0.05, **p<0.01, ***p<0.001, ****p<0.0001). **g**) Kymographs depicting active transport of MYO3A and PCDH15-CD2 along a representative filopodium. Time elapsed from the first frame to the last was 12 seconds. The green and magenta fluorescence puncta correspond to clusters of MYO3A and PCDH15-CD2 co-translocating along the filipodium. Scale bars = 2 μm.

To confirm that the FL PCDH15-CD2 isoforms can also be transported by MYO3A to filopodia tips, we performed similar experiments with FL PCDH15-CD1, -CD2 and -CD3 (Fig. 2d-f). We observed that FL PDH15-CD2, but not -CD1 or -CD3, was enriched at the tips of MYO3A filopodia (Fig. 2e). In fact, we observed a higher tip to shaft ratio of FL PCDH15-CD2 when compared to cyto PCDH15-CD2, while the expression of MYO3A did not alter the cellular localization of the -CD1 or -CD3 isoforms. However, the increase in the MYO3A filopodia tip to shaft ratio with FL PCDH15-CD2 (Fig. 2f) was not as striking as with the cyto PCDH15-CD2 and MYO3A. Additionally, live-cell imaging in COS7 cell expressing fluorescently tagged MYO3A and cyto-PCDH15-CD2 shows that the localization and enrichment of PCDH15-CD2 at filopodia tips is the result of active transport by MYO3A. Filopodia of COS7 cells transfected with GFP-MYO3A and mCherry-cyto-CD2 showed dynamic localization of both proteins at filopodia tips from the early steps of their initiation and elongation. Kymographs of the filopodia also showed that the two proteins leave the filopodia tips and co-translocate towards the filopodia base (Fig. 2g). This retrograde motion is due to clusters of the myosin and its cargo riding the reward treadmilling of the actin that makes up the filopodia core ^38^, indicating that MYO3A and PCDH15-CD2 maintain interaction as they translocate along actin.

### Espin-1 and espin-like differentially regulate the transport of PCDH15-CD2 by class III myosins

The class III myosin cargoes espin-1 or espin-like bind to the tail homology domain I (THDI) of MYO3A and MYO3B to differentially regulate molecular transport and control filopodia growth and are essential for normal hearing^29^. Therefore, we next examine whether expression of espin-1 and espin-like alters the transport of PCDH15-CD2 FL to the filopodia tips.

Cells expressing MYO3A, PCDH15-CD2 and espin-1 presented the characteristic long filopodia of cells expressing a class III myosin and espin-1^48^. However, while cells expressing MYO3A and PCDH15-CD2 FL have these two proteins enriched at their filopodia tips, PCDH15-CD2 FL is no longer enriched at the filopodia tips in cells expressing MYO3A and espin-1 (Fig. 3a and c), suggesting that espin-1 prevents the transport of PCDH15-CD2 by MYO3A. We next analyzed the transport of PCDH15-CD2 by MYO3A in the presence of the espin-1 paralog, espin-like. Importantly, we focus our study on cells expressing low levels of espin-like, since high expression levels inhibit MYO3A-dependent filopodia formation^29^. In contrast to that observed with espin-1, PCDH15-CD2 FL was enriched at the filopodia tips of COS7 cells expressing MYO3A and espin-like (Fig. 3a) although PCDH15-CD2 FL was less enriched at the MYO3A filopodia tips in the presence of espin-like than in the absence (Fig. 3c), suggesting that MYO3A can transport espin-like and PCDH15-CD2 simultaneously. Quantification of MYO3A enrichment at the filopodia tips in absence or presence of espin-like revealed that espin-like did not affect MYO3A enrichment (Fig. 3d). These data indicate that the presence of espin-like does not affect the transport of PCDH15-CD2 by MYO3A and reveal an intricate regulatory role for MYO3A cargos in the transport of PCDH15-CD2.

**Figure 3:**
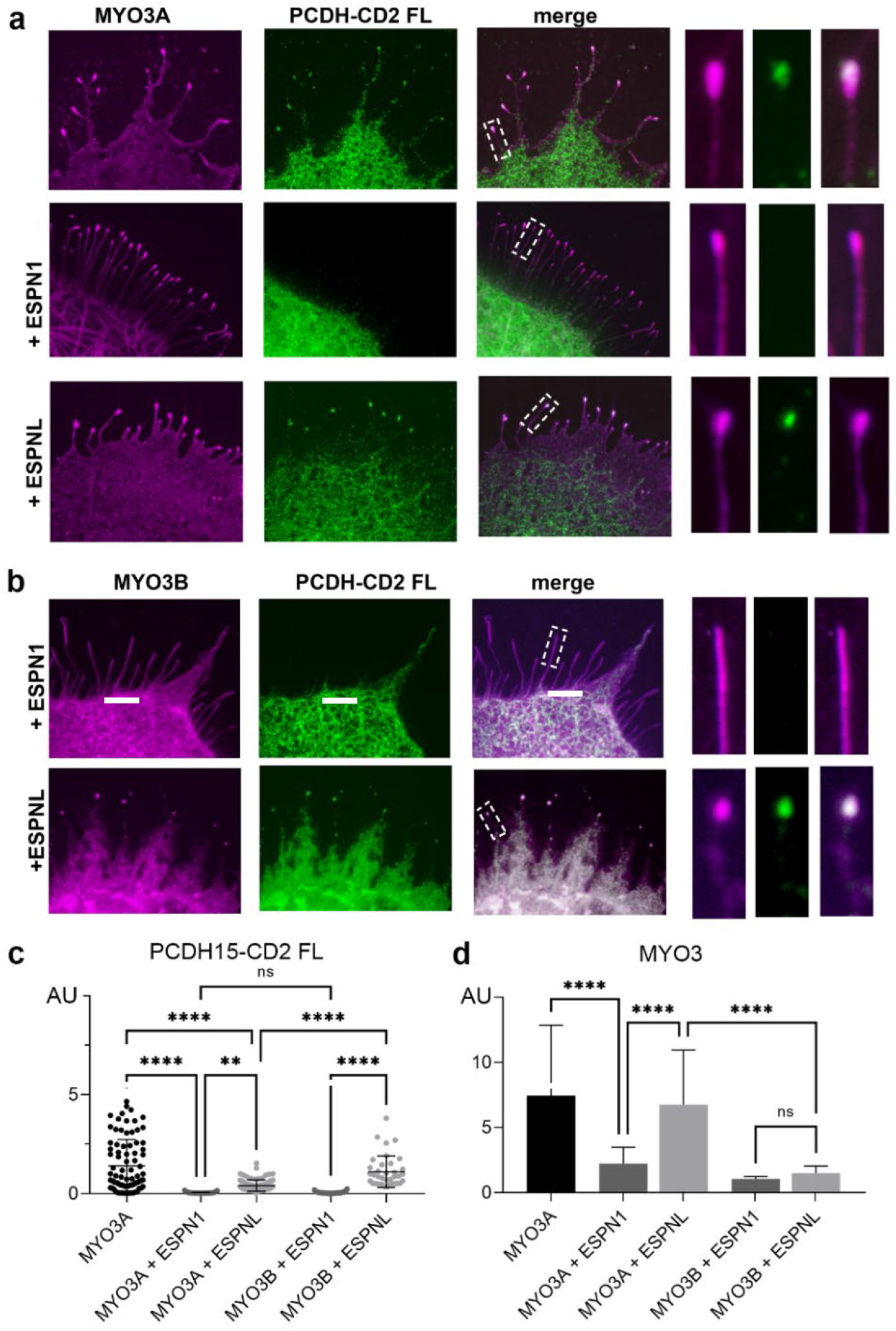
Espin-1 and espin-like differentially regulate the transport of PCDH15-CD2 by class III myosins. **a**) Confocal images of COS7 cells expressing GFP-MYO3A (magenta) and PCDH15-CD2 FL (green) in the absence or presence of espin-1 (ESPN1) or espin-like (ESPNL) (**b**). Confocal images of COS7 cells expressing GFP-MYO3B (magenta), PCDH15-CD2 FL (green) and espin-1 or espin-like. Filopodia zoom in images are shown at the right. **c**) PCDH15-CD2 enrichment at the filopodia tips in the conditions shown in a and b. Mean ± SD is shown. Each dot represents one filopodia. **d)** MYO3A relative enrichment at filopodia tips for each of the conditions examined. One-way ANOVA analysis was performed with Dunnett’s multiple comparisons test. Level of significance was determined based on the p value (n.s p>0.05, *p<0.05, **p<0.01, ***p<0.001, ****p<0.0001). Scale bar = 2 μm.

The shorter myosin class III isoform, MYO3B, lacks the additional actin-binding domain that MYO3A presents on its tail (3THDII) (Fig. 1c), which prevents this myosin from translocating along actin and concentrating at the tips of filopodia by itself ^7,39^ Consequently, to move along the actin filaments and reach the tips of stereocilia or filopodia, MYO3B requires the actin-binding activity of espin-1^7^ or espin-like^29^. We thus examined whether MYO3B transports PCDH15-CD2 FL to the filopodia tips when co-expressed with espin-1 or espin-like in COS7 cells. We first observed that MYO3B enrichment at the filopodia tips is less efficient than MYO3A, as previously reported^7^, and that expression of espin-1 or espin-like did not alter MYO3B enrichment (Fig. 3d). Like that observed with MYO3A, we found that PCDH15-CD2 FL was enriched at the filopodia tips of cells expressing MYO3B and espin-like but not espin-1 (Fig. 3b and c), indicating that espin-1 and espin-like differentially regulate the transport of PCDH15-CD2 by MYO3A and MYO3B.

### MYO7A transports PCDH15-CD3 to the tips of filopodia

Immunoprecipitation experiments have shown that the tail domain of MYO7A interacts with the cytoplasmic domain of PCDH15 cloned from mouse brain tissue^40^, suggesting that these proteins could cooperate to regulate hair cell bundle development and function. Contrary to the other myosins tested in this study, class VII myosins maintain a cytoplasmic non-motile conformation and do not form filopodia by themselves ^35^. Therefore, to test the role of MYO7A in the transport of the three PCDH15 isoforms, we used a chimeric MYO7A/MYO10 myosin containing the tail cargo-binding domain of MYO7A and the motor domain of MYO10 (Fig. 1c), which is sufficient to translocate on actin and induce filopodia formation ^35,41,42^

Cell expressing MYO7A/MYO10 and PCDH15-CD3 show co-enrichment of the two proteins at the tip of filopodia (Fig. 4a-b). However, we did not observe enrichment of PCDH15-CD1 nor PCDH15-CD2 in cells expressing MYO7A/MYO10, suggesting that MYO7A tail specifically binds to PCDH15-CD3. Furthermore, we did not observe any difference in the enrichment of MYO7A/MYO10 at the tips of filopodia when expressed in combination with PCDH15-CD1, -CD2, or -CD3 (Fig. 4c), suggesting that the PCDH15-CD3 cargo is not influencing MYO7A/MYO10 motor activity. To confirm that PCDH15-CD3 transport is dependent on the MYO7A tail, and not the MYO10 portion of the chimeric MYO7A/MYO10, we performed similar experiments using MYO10 heavy meromyosin (HMM), consisting of the head, neck, and α-helical region of MYO10. While MYO10 was equally enriched at the tips of filopodia of COS7 cell it was not accompanied with tip enrichment of any of the PCDH15 isoforms including the CD3 (Fig. 4d-e). These data suggest that MYO7A tail is required for the transport of PCDH15-CD3.

**Figure 4:**
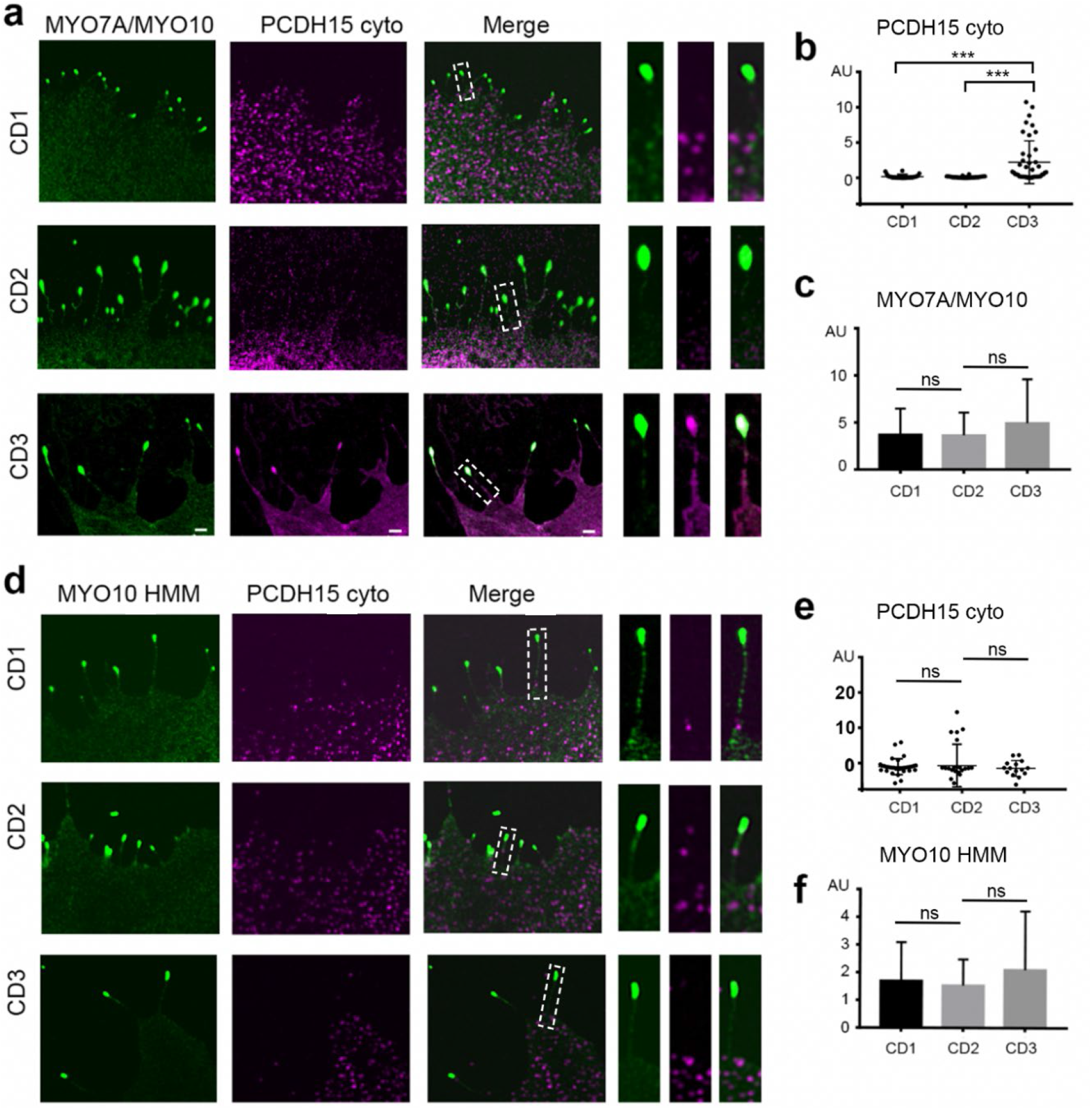
MYO7A transports PCDH15-CD3 to the tips of filopodia. **a**) Confocal images of COS7 cells expressing the chimeric MYO7A/MYO10 protein (green) and cyto PCDH15-CD1, -CD2 or -CD3 (magenta). The right panels show a close-up view of a representative filopodia. **b**) PCDH15-CD3 cyto exhibits significantly higher tip to cell ratio than PCDH15-CD1 or PCDH15-CD2 when co-expressed with MYO7A/MYO10. **c)** MYO7A/MYO10 exhibit a similar enrichment at the filopodia tips when co-transfected with FL PCDH15-CD1, -CD2 or -CD3. **d**) Confocal images of COS7 cells expressing the MYO10 (green) and cyto PCDH15-CD1, -CD2 or -CD3 (magenta). The right panels show a close-up view of a representative filopodia. **e**) We did not observe a significantly change in the tip to cell ratio of any of the PCDH5 isoform tested. **f)** MYO10 HMM exhibited a similar enrichment at the filopodia tips when co-transfected with FL PCDH15-CD1, -CD2 or -CD3. Each dot in b and e represents one filopodia. One-way ANOVA analysis was performed with Dunnett’s multiple comparisons test. Level of significance was determined based on the p value (n.s p>0.05, *p<0.05, **p<0.01, ***p<0.001, ****p<0.0001). Scale bar = 2 μm.

Interestingly, MYO7A cargoes have been shown to promote the dimerization and filopodia formation activity of MYO7A^42^. Since our data suggest that PCDH15-CD3 might be a MYO7A cargo, we examined whether PCDH15-CD3 can promote filopodia formation by wild type MYO7A. COS7 cells expressing PCDH15-CD3 and MYO7A showed a diffuse intracellular localization of MYO7A and do not present filopodia, indicating that PCDH15-CD3 does not activate MYO7A to move along actin structures as a cargo transporting motor (Supplementary Figure 4).

### MYO15A, a principal cargo transport in stereocilia, does not transport PCDH15 to the tips of filopodia

MYO15A transports whirlin and eps8 to the stereocilia tips and mutation in these proteins results in shortening of stereocilia and hearing loss ^8,43,44^. Using the same co-expression assay, we tested the potential role of MYO15A in the transport of PCDH15 isoforms. We observed that while MYO15A induces filopodia formation and is enriched at the filopodia tips of COS7 cells (Fig. 5a), it did not promote the enrichment of PCDH15-CD1, -CD2 nor -CD3 at the filopodia tips (Fig. 5b). Therefore, our data does not support a direct role for MYO15A in the transport of any of the three main PCDH15 isoforms in stereocilia.

**Figure 5:**
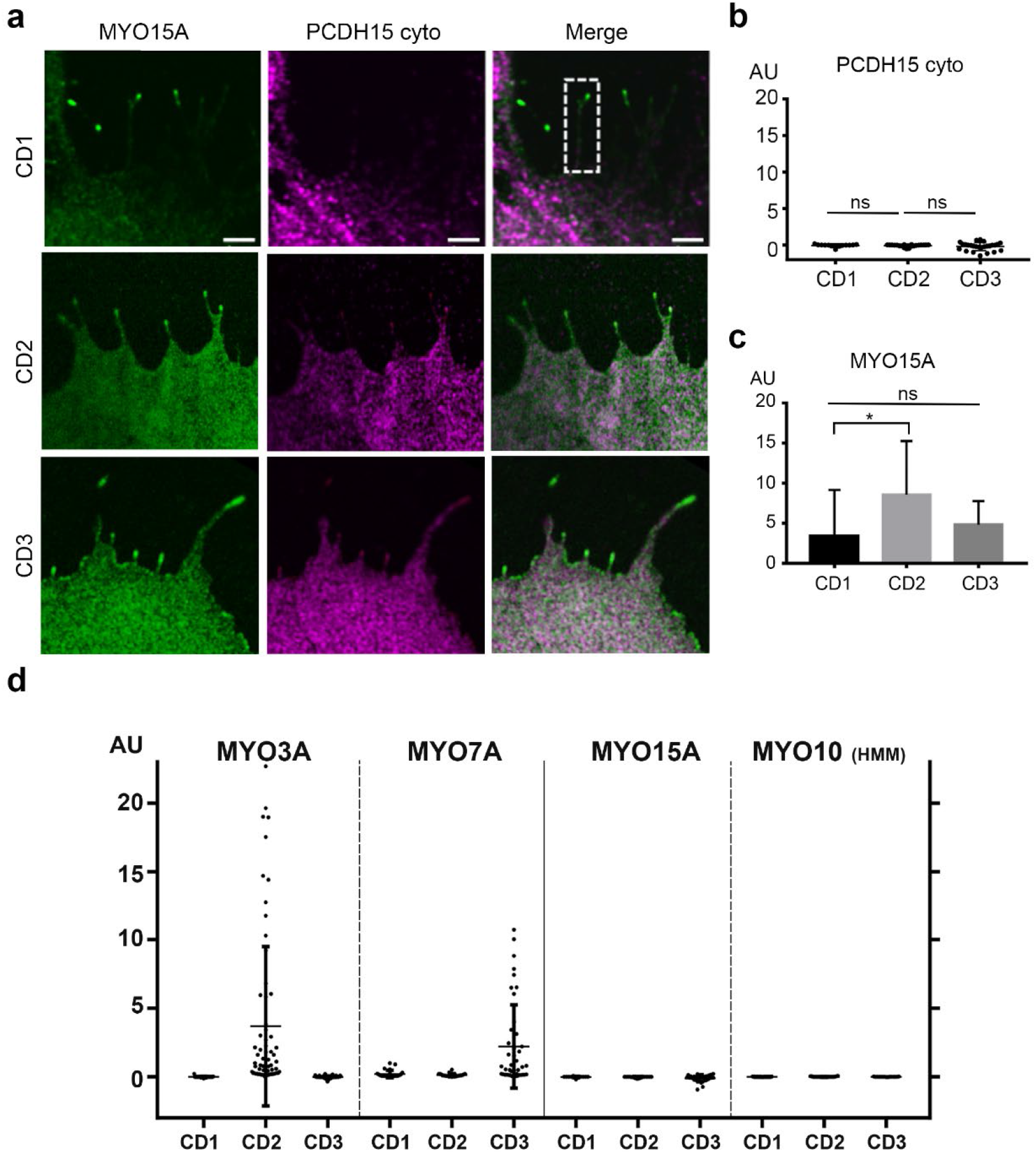
MYO15A does not transport PCDH15 to the tips of filopodia. **a**) Confocal images of COS7 cells expressing MYO15A (green) and cyto PCDH15-CD1, -CD2 or -CD3 (magenta). **b**) Any of the PCDH15 isoforms showed a significantly tip to cell ratio when co-expressed with MYO15A. **c)** MYO15A exhibit a slightly higher enrichment at the filopodia tips when co-transfected with cyto PCDH15-CD2 when compared to -CD1 or -CD3. **d**) Summary of the tip to shaft ratio of PCDH15 isoforms when expressed with MYO3A, MYO7A/MYO10, MYO15A and MYO10. Each dot in b and d represents one filopodia. One-way ANOVA analysis was performed with Dunnett’s multiple comparisons test. Level of significance was determined based on the p value (n.s p>0.05, *p<0.05, **p<0.01, ***p<0.001, ****p<0.0001). Scale bar = 2μm

## DISCUSSION

The mechanisms of transport and assembly of molecular components of the MET apparatus at stereocilia tips remains a key question in hair cell biology. The characterization of direct interactions and individual roles for each molecular MET component has been challenging due to the molecular crowding at stereocilia tips, the mutual influence that each protein can exert on each other, and the compensatory roles reported for several molecular components. In this work, we took a heterologous expression approach to dissect the role myosins commonly detected in stereocilia may have in the selective transport of PCDH15 isoforms. With this approach, we demonstrate that MYO3A selectively transports PCDH15-CD2 (Fig. 2) while MYO7A transports PCDH15-CD3 (Fig. 4). Surprisingly, MYO15A, which is another key stereocilia cargo transport myosin and localizes at stereocilia tips did not show interactions with any of the three PCDH15 isoforms (Fig. 5). However, our results are consistent with the reported preservation of MET in the shaker-2 mouse model lacking MYO15A^45^.

Interestingly, none of the myosins tested showed effective transport of the PCDH15-CD1 isoform. Senften and colleagues^40^ reported that PCDH15 from mouse brain binds to the SRC Homology 3 (SH3) domain of MYO7A and recruits MYO7A to the cell membrane in cultured cells. While the specific PCDH15 isoform used in this study was not specified, its sequence length corresponds to the CD1 isoform. It is possible that other factors or adaptor proteins may be required for the interactions of PCDH15-CD1 with MYO7A or other stereocilia myosins. Cargo or adaptor-dependent regulation of myosin transport has been established in the literature ^46^ and is also evident in our experiments where espin1 interfered with the MYO3A-dependent enrichment of PCDH15-CD2 at filopodia tips (Fig. 3).

Myosins and espins are directly involved in regulation of actin filament formation and stability including in the differential regulation of stereocilia length. Espin-1 and espin-like bind to the THDI of MYO3A and MYO3B and can mutually influence myosin activity and actin polymerization^30^. Differential expression of these proteins correlates with differential lengths of stereocilia in each bundle and across different hair cells along the organ of Corti or in the different inner ear sensory epithelia^29,47^. The dynamic localization of PCDH15-CD2 at filopodia tips when co-transfected with MYO3A indicate that MYO3A specifically interacts with and transports PCDH15-CD2. PCDH15-CD2 transport by MYO3A and MYO3B in the presence of espin-like suggest that PCDH15-CD2 may interact with a common domain in these class III myosins. nhibition of PCDH15-CD2 transport by espin-1 suggest that these two MYO3A cargoes could compete for a common MYO3A binding site. However, in this scenario, we would have expected espin-like to also inhibit PDH15-CD2 transport since espin-1 and espin-like bind to the THD1 domain of MYO3A and MYO3B in a conserved way^48^. Therefore, a more complex mechanism may be in place by which espin1 and espin-like can differentially regulate PCDH15-CD2 transport and enrichment at the tips of actin protrusions. Espin 1 and espin-like are known to influence the length and structure of filopodia and stereocilia ^7,29,47,49^ It is also likely that the transport and enrichment of cargoes depend on the length and number of actin tracks in an actin protrusion. Consequently, the transport of actin regulatory proteins by MYO3A and 3B may influence the transport of PCDH15-CD2 in multiple ways and modulate the relationship between stereocilia length and MET.

Our results highlight the intricate nature of the mechanisms of molecular cargo transport and assembly of the MET complex that takes place during development, maturation, life-long function, and likely in the degeneration of the stereocilia bundles during aging. Several isoforms of PCDH15 differing in their cytoplasmic domains are expressed in the hair cell bundle, suggesting that alternative splicing regulates PCDH15 localization and function. The main three PCDH15-CD1, PCDH15-CD2 and PCDH15-CD3 isoforms can compensate for each other in the radial-link formation and the tip-link formation of immature hair cells^25,26^ (Fig. 1a). However, PCDH15-CD2 has an essential role in the formation of the tip-link in mature hair cells. Another essential component of the MET channel complex is the membrane protein TMIE, which has been shown to specifically interact with PCDH15-CD2^28,50^. Interestingly, appearance and expression levels of MYO3A at stereocilia tips during development are correlated with onset and maturation of MET^51^. Taken together these data suggest that MYO3A and MYO3B play a role in the assembly and maturation of the MET complex. Given that MYO3A and MYO3B are intra- and inter-molecularly regulated by their kinase domain and the presence several calmodulin binding sites, they can also potentially exhibit Ca^2+^-dependent regulation^52–54^. Therefore, we cannot rule out their participation in the MET adaption processes. Consistent with this, a recent report provides good evidence that MET adaptation takes place at or near the channel complex^55^.

While PCDH15 has not been shown to directly affect actin regulation, its proper targeting to the stereocilia tip and participation in the tip link and MET is required for normal length regulation of stereocilia as it has been shown in various experimental conditions and mouse models^3,4,56^. Furthermore, the selective transport of PCDH15-CD2 and PCDH15-CD3 by MYO3A and MYO7A could regulate the composition and properties of the MET channel influencing the transport of other essential MET complex components know to interact with PCDH15^50,57,58^. Interestingly, all the myosins and the cargoes considered in this study have been directly involved in multiple syndromic and non-syndromic forms of hearing loss and vestibular dysfunction ^30,43,59–63^. Our results showing the intricate relationships between these molecular components provide new insights into potential common molecular and structural mechanisms underlying loss of function.

Our observations in the reductionist heterologous expression system allowed us to identify specific interactions that would otherwise be concealed in the complex and interconnected context of stereocilia bundles in mouse models. However, additional work is necessary to elucidate how PCDH15-CD1 and other component of the MET channel complex are transported, and how the myosin and cargo systems are regulated and the extent of their cooperativity and complementarity in normal and affected hair cells during development, homeostasis, and aging.

## MATERIAL AND METHODS

### Expression plasmids

The full length of human PCDH15-CD1-1 (Q99PJ1, uniport) cloned in pcDNA3.1 was obtained from OriGene. PCDH15-CD2 (Q99PJ1-10, uniport) and PCDH15-CD3-1 (Q99PJ1-18, uniport) were cloned from genomic mouse DNA. The cytoplasmic domains of the PCDH15 isoforms corresponding to residues 1403-1443 of PCDH15-CD1, 1410-1790 of PCDH15-CD2, and 1403-1682 of PCH15-CD3, were PCR amplified from the full-length cDNAs and cloned into the NheI and BamHI or into the HindIII and BamHI restriction sites of the pEGFP-C2 vectors (Clontech), resulting in untagged soluble proteins or proteins tagged with a GFP fused to its N-terminus, respectively. For live imaging experiments, the PCDH15-CD2 isoform cytoplasmic domain was cloned using the HindIII and BamHI restriction sites of the pmCherry-C2 vector (Clontech), resulting in soluble protein tagged with cherry at its N-terminus.

The human MYO3A and mouse MYO3B constructs used in our study lack the N-terminal kinase domain of class III myosins (MYO3A-ΔK and MYO3B-ΔK) and thus localize more efficiently to the stereocilia tips than the full-length protein. The kinase domain has been shown to downregulate motor activity leading to a slower Myosin3A protein ^33^. The espin-1, espin-like, GFP-MYO3A, GFP-MYO3B, and mCherry-MYO3B cDNAs were previously prepared and used in our laboratory and described ^7,29,47^

### Antibodies

Affinity purified rabbit polyclonal anti-mouse PCDH15 antibody (COVANCE) generated against the N-terminus domain of PCDH15 conserved in all the PCDH15 isoform was used to detect all the PCDH15 isoforms (pan-PCDH15 antibody, PB808). Affinity purified rabbit polyclonal anti-mouse PCDH15 antibodies (COVANCE) generated against the variable intracellular cytoplasmic domain (CD) was used to specifically detect the PCDH15 isoforms PCDH15-CD1, PCDH15-CD2, and PCDH15-CD3 (PB246, PB860, and PB814, respectively) (Table 1). The four custom-made antibodies used in this manuscript were validated in transfected COS7 cells expressing the three PCDH15 isoforms as shown in the Supplementary Fig. 2 and explained in the following confocal microscopy analysis section.

### Confocal microscopy imaging

COS7 cells (ATCC, CRL-1651) were grown and maintained in Dulbecco’s Modified Eagle Medium (DMEM) supplemented with 10% fetal bovine serum (FBS) at 37°C and 5% CO2 in a cell incubator. Cells were trypsinized and plated on glass coverslips at a 50% confluency, and 18 h later, they were transfected with 1μg of cDNA using the Lipofectamine transfection reagent (Invitrogen) per manufacturer’s instructions. Cell media wash changed 18 h after transfection to remove the residual Lipofectamine and cells were maintained in the cell incubator during 24 h for protein expression. Samples were then fixed for 20 min in 2% paraformaldehyde in Phosphate Buffered Saline (PBS), permeabilized and counterstained for actin with a 1:100 dilution of CF-405M Phalloidin (Biotium) in 0.5% Triton X-100 in PBS for 30 min. After removing TritonX-100 and phalloidin with 2-3 PBS washes, samples were mounted on glass slides and imaged in a Nikon microscope equipped with a Yokogawa spinning disk confocal unit.

When expressing the untagged PCDH15 isoforms, cells were permeabilized 0.5% Triton X-100 in PBS for 30 min and the excess Triton X-100 was were removed with 2-3 PBS washes, and 4% Bovine Serum Albumin (BSA) in PBS was added to the cells and incubated for 1h. Primary rabbit polyclonal antibodies against PCHD15 (PB808, PB246, PB860, or PB814) at a 1:500 dilution in PBS with 4% BSA was added and incubated with the cells for 1h at RT. After 2-3 washes with PBS to remove unbound antibody, secondary Alexa flour 564 or Alexa flour 643 anti-rabbit (Life Technology) at a 1:2000 dilution in PBS with 4% BSA was added and incubated for 30 min together with CF405M-Phalloidin (Biotium) at a 1:100 dilution to label F-actin. Cells were then washed with PBS several times and mounted using Celvol 205 mounting media in microscopy slides for confocal imaging. Imaging was performed in a Nikon microscope equipped with a Yokogawa spinning disk confocal unit.

### Quantification of fluorescently tagged and immunolabeled proteins along the filopodia

Microscopy data processing, analysis, and estimation of the relative pixel intensity of fluorescently tagged proteins along the filopodia were done in ImageJ ^64^. To quantify the enrichment at the filopodia tips, we measured the filopodia tip to shaft fluorescence intensity ratio. To do this, we used the line tool from ImageJ to draw three lines of one pixel in width; one along the filopodia tip, other one along the filipodia shaft, and a third one in a region outside filopodia. The line drew near but outside the filopodia was considered as background and subtracted from the fluoresce measurements. We calculated the ratio of tip vs shaft or estimated the enrichment at the filopodia tips by subtracting the fluorescent intensity at the filopodia shaft from the one at the tip. To measure the enrichment at the filopodia, we measure the filopodia tips to cell fluorescence intensity ratio. The data was imported into the GraphPad Prism 8 software for the generation of the graphs and statistical analysis.

### Immunolabeling of mammalian hair cells

All mice used in this work were handled following the NIH Guidelines for Animal Care and approved by the NIDCD Animal Protocol Committee (Protocol #1215). Cochlear tissue from wild type 10 days old mice or vestibular tissue from adult guinea pig was rapidly isolated in Leibovitz L-15 medium and fixed in 4% formaldehyde in HBSS buffer for 20 min and then washed with phosphate-buffered saline (PBS). Samples were then permeabilized with 0.5% Triton X-100 in PBS for 20 min before blocking with bovine serum albumin (BSA). After blocking for 1 h with 1% BSA, 3% normal goat serum in PBS, organs were incubated overnight at 4°C with primary antibodies in blocking buffer. Custom-made pan PCDH5 antibody PB811 or a mixture of the pan PCDH15 containing PB811 and PB808 antibodies were used to detect all the PCDH15 isoforms. Organs were then washed 3 × 5 min with PBS and incubated with secondary antibodies (Alexa 546-conjugated goat anti-rabbit IgG) containing Alexa Fluor 405 phalloidin (to label stereocilia actin) for 30 min, before washing five times in PBS and mounting between a microscope slide and cover slip. Samples were viewed in a Nikon inverted fluorescence microscope, fitted with a spinning disk confocal scan head (Yokogawa, CSU-22), 100× Apo 1.49 numerical aperture objective, and an EM-CCD (Andor 888 or 897) camera. Nikon NIS-Elements imaging software was utilized for image acquisition and analysis.

## DATA AVAILABILITY

The datasets generated and/or analyzed during the current study are available from the corresponding author on reasonable request.

## ACKNOLEDGMENTS

We are greatly thankful to the Robert Wenthold postdoctoral fellowship and the National Institute on Deafness and other Communication Disorders Intramural Research Program, National Institutes of Health (NIDCD-NIH) for individual support to A.B. Work was supported by NIDCD-NIH-IRP funds Z01-DC000002. We would like to thank Dr. M’hamed Grati, NIDCD, NIH for the cDNA encoding the chimeric MYO7A/MYO10 and Dr. Melanie Barzik, NIDCD, NIH for critical review of the manuscript.

## AUTHOR CONTRIBUTIONS

A.B., M.Y., and R.C. performed the experiments. K.K. and A.B. generated the cDNA constructs. A.B., M.Y. and B.K. conducted the data collection. B.K. and A.B. constructed the concept. M.Y performed the image analysis. A.B. and B. K. wrote the manuscript and all the authors proofread and ensured the general quality of the manuscript.

## COMPETING INTEREST

The authors declare no conflict of interest.

## SUPPEMENTARY INFORMATION

**Supplementary Figure 1:**
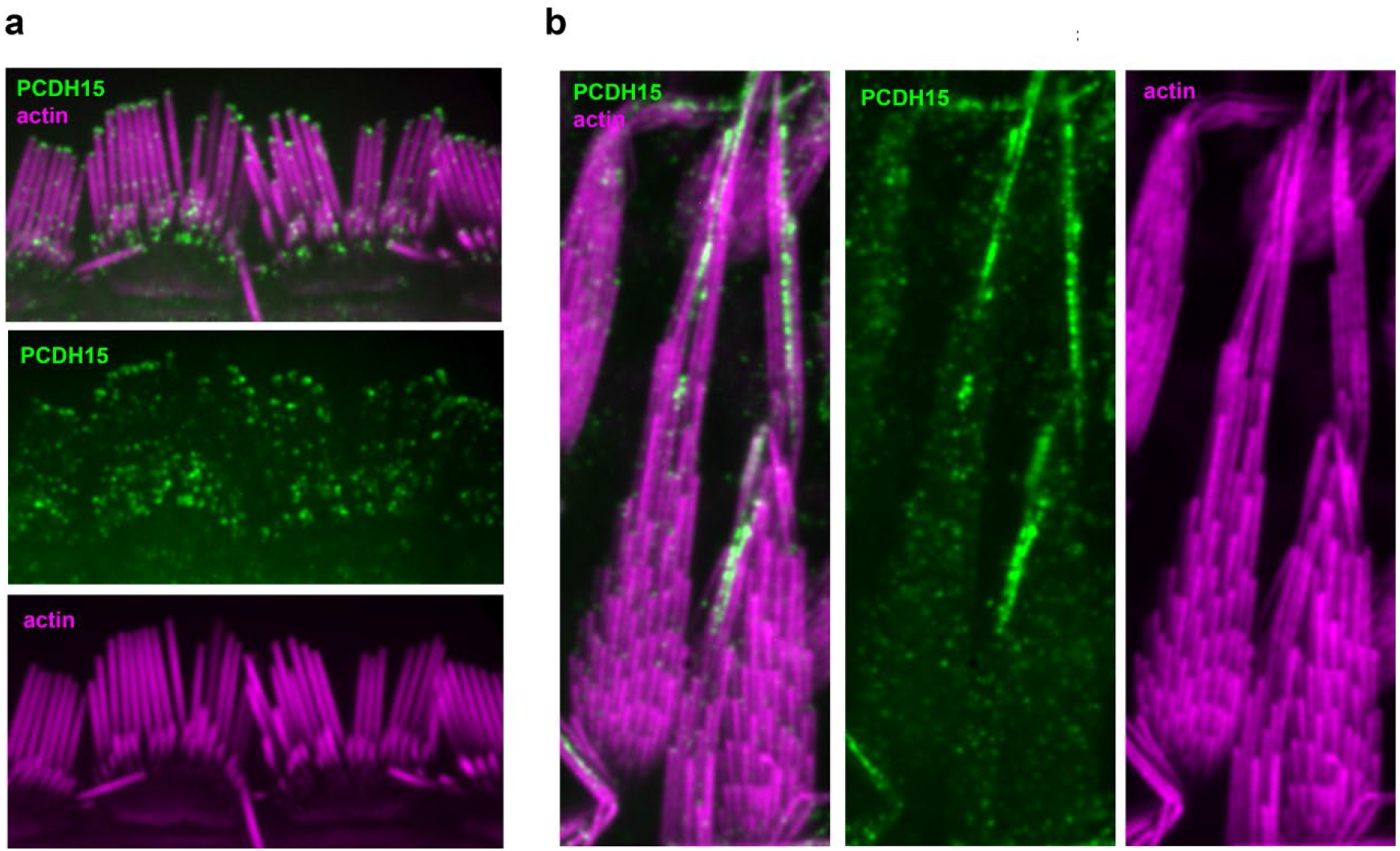
PDH15 localization in mammalian auditory and vestibular hair cells. **a)** PCDH15 localizes to the tips of the stereocilia in 10 days old murine inner hair cells (IHC) as showed by immunohistochemistry with a pan PCDH5 antibody PB811 (green). **b**) PCDH15 (green) is accumulated at the kinociliary links in vestibular hair cells from adult guinea pig as showed by immunohistochemistry with a pan PCDH5 antibody. Tissue explants were counterstained with phalloidin to label F-actin and visualize the hair cell stereocilia (magenta). Scale bar = 5 μm.

**Supplementary Figure 2:**
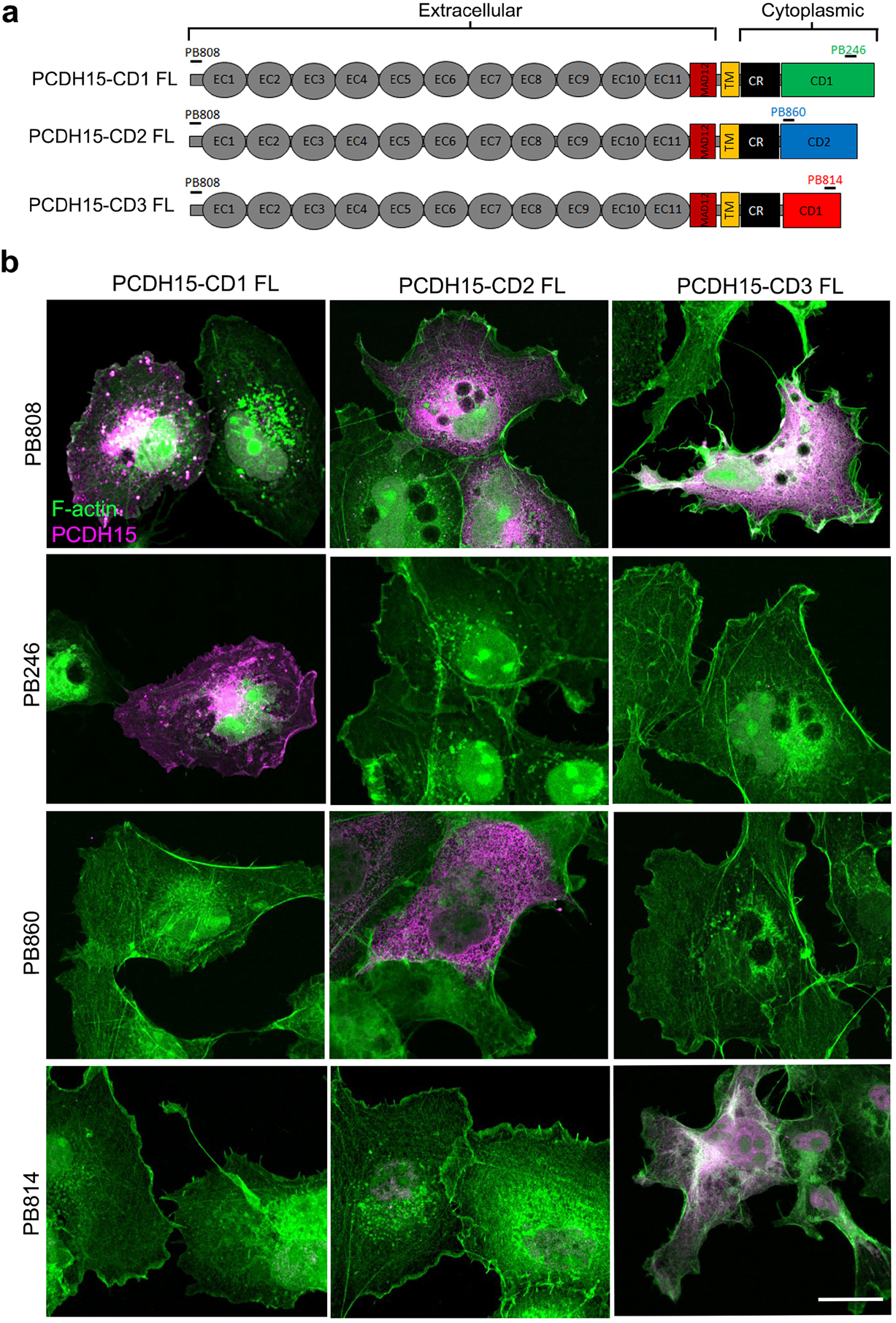
Validation of the anti PCDH15 antibodies and the Full-length PCDH15 constructs. **a)** Diagram of the PCDH15-CD1, -CD2 and -CD3 constructs used in this study and localization of the peptides used to generate the PCDH15 antibodies PB808, PB246, PB860 and PB814. **b**) COS7 cells expressing any of the three PCDH15 isoforms (PCDH15-CD1, -CD2 or -CD3) were labeled with the pan-PDH15 antibody PB808, while only cells expressing the specific PCDH15 isoform were labeled with the isoform-specific anti PCDH15 antibody (magenta). Cells were counterstained with phalloidin to label F-actin (green). Scale bar = 15 μm.

**Supplementary Figure 3:**
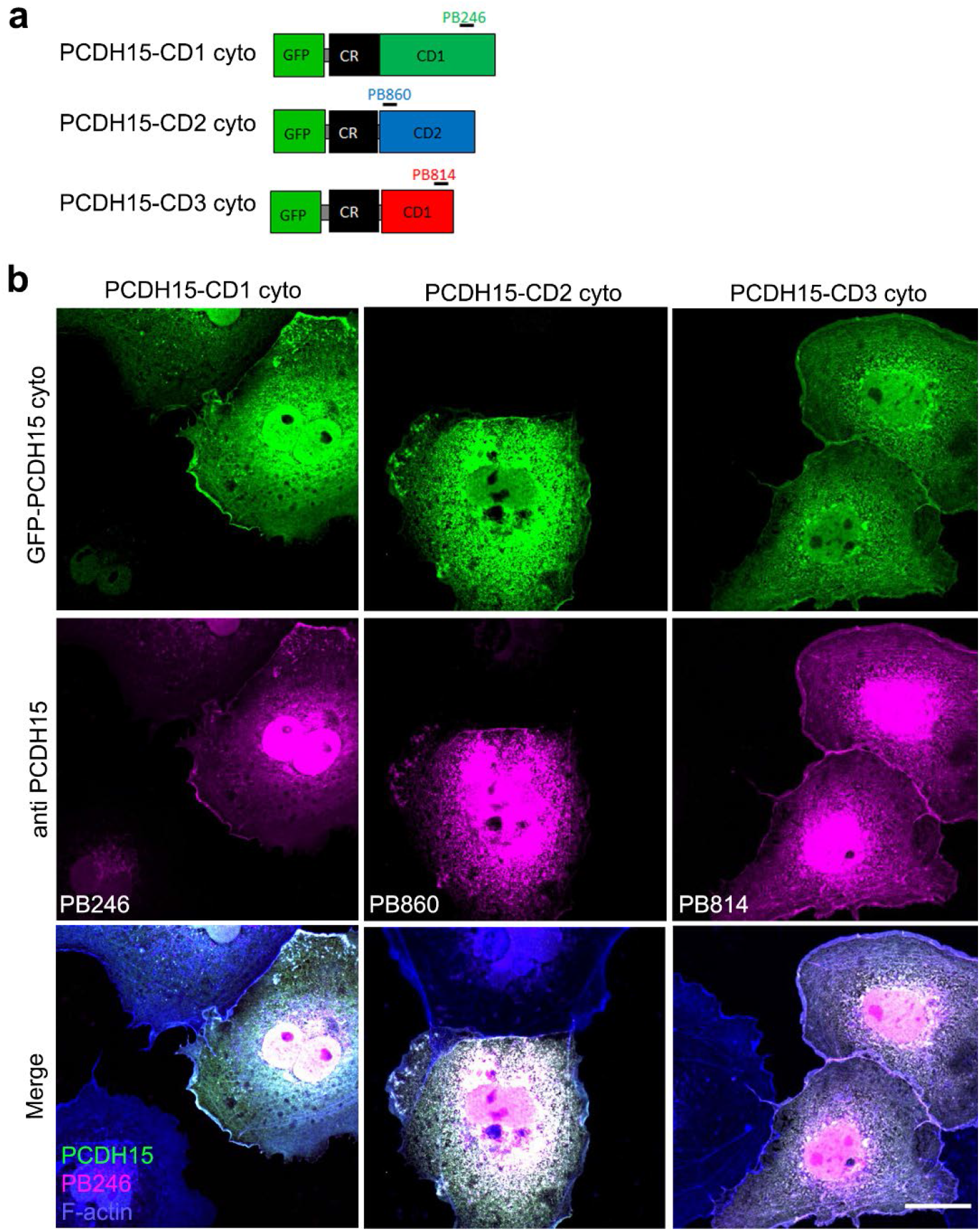
Validation of the anti PCDH15 antibodies and the cyto PCDH15 constructs. **a)** Diagram of the cyto GFP-PCDH15-CD1, -CD2 and -CD3 constructs used in this study and localization of the peptides used to generate the isoform-specific PCDH15 antibodies PB246, PB860 and PB814. **b**) COS7 cells expressing the specific GFP tagged PCDH15 isoform (green) were also labeled with the corresponding isoform specific anti PCDH15 antibody (magenta). Cells were counterstained with phalloidin to label F-actin (blue). Scale bar = 15 μm.

**Supplementary Figure 4:**
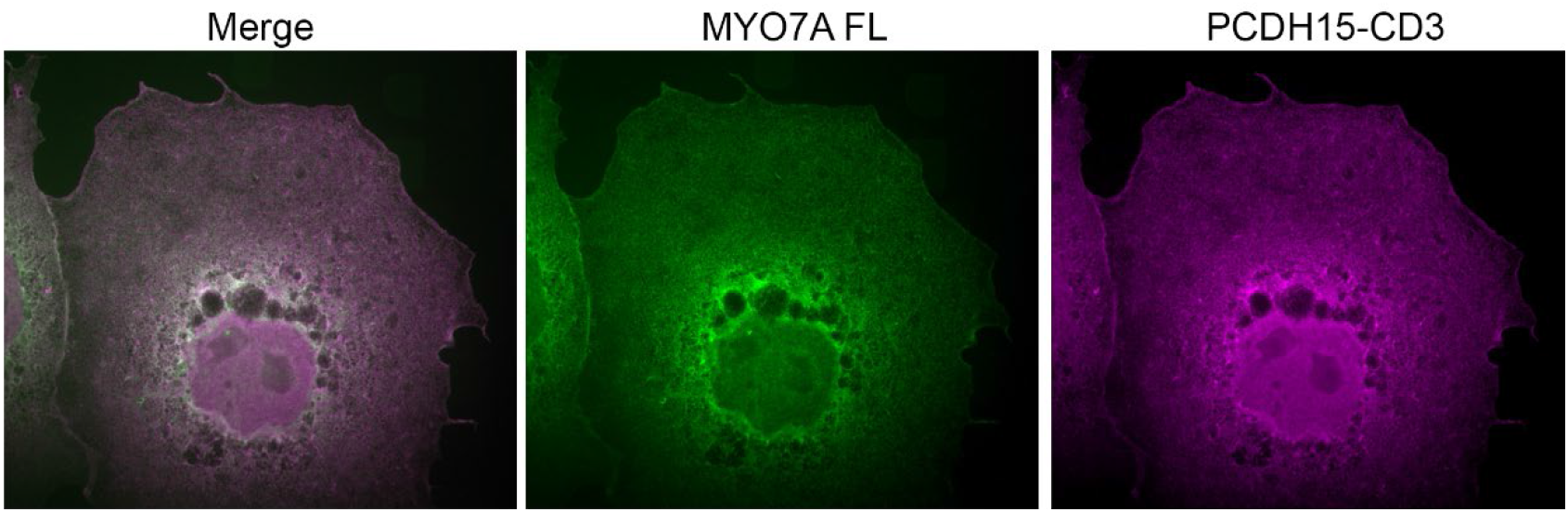
Myosin 7A does not form filopodia when expressed in COS7 cells. COS7 cell expressing MYO7A full length (MYO7AFL) and PCDH15-CD3 shows that PCDH15-CD3 does not activate MYO7A. Scale bar = 15 μm.

## Notes

### Competing Interest Statement

The authors have declared no competing interest.

## REFERENCES

1 Schwander, M., Kachar, B. & Muller, U. Review series: The cell biology of hearing. J Cell Biol 190, 9–20, doi:10.1083/jcb.201001138 (2010).

2 Beurg, M., Fettiplace, R., Nam, J. H. & Ricci, A. J. Localization of inner hair cell mechanotransducer channels using high-speed calcium imaging. Nat Neurosci 12, 553–558, doi:10.1038/nn.2295 (2009).

3 Krey, J. F. et al. Mechanotransduction-Dependent Control of Stereocilia Dimensions and Row Identity in Inner Hair Cells. Curr Biol 30, 442–454 e447, doi:10.1016/j.cub.2019.11.076 (2020).

4 Velez-Ortega, A. C., Freeman, M. J., Indzhykulian, A. A., Grossheim, J. M. & Frolenkov, G. I. Mechanotransduction current is essential for stability of the transducing stereocilia in mammalian auditory hair cells. Elife 6, doi:10.7554/eLife.24661 (2017).

5 McGrath, J. et al. Actin at stereocilia tips is regulated by mechanotransduction and ADF/cofilin. Curr Biol 31, 1141–1153 e1147, doi:10.1016/j.cub.2020.12.006 (2021).

6 Hartman, M. A., Finan, D., Sivaramakrishnan, S. & Spudich, J. A. Principles of unconventional myosin function and targeting. Annu Rev Cell Dev Biol 27, 133–155, doi:10.1146/annurev-cellbio-100809-151502 (2011).

7 Merritt, R. C. et al. Myosin IIIB uses an actin-binding motif in its espin-1 cargo to reach the tips of actin protrusions. Curr Biol 22, 320–325, doi:10.1016/j.cub.2011.12.053 (2012).

8 Manor, U. et al. Regulation of stereocilia length by myosin XVa and whirlin depends on the actin-regulatory protein Eps8. Curr Biol 21, 167–172, doi:10.1016/j.cub.2010.12.046 (2011).

9 Belyantseva, I. A. et al. Myosin-XVa is required for tip localization of whirlin and differential elongation of hair-cell stereocilia. Nat Cell Biol 7, 148–156, doi:10.1038/ncb1219 (2005).

10 Kros, C. J. et al. Reduced climbing and increased slipping adaptation in cochlear hair cells of mice with Myo7a mutations. Nat Neurosci 5, 41–47, doi:10.1038/nn784 (2002).

11 Walsh, V. L. et al. A mouse model for human hearing loss DFNB30 due to loss of function of myosin IIIA. Mamm Genome 22, 170–177, doi:10.1007/s00335-010-9310-6 (2011).

12 Qu, R. et al. Identification of a novel homozygous mutation in MYO3A in a Chinese family with DFNB30 non-syndromic hearing impairment. Int J Pediatr Otorhinolaryngol 84, 43–47, doi:10.1016/j.ijporl.2016.02.036 (2016).

13 Dantas, V. G. L. et al. Characterization of a novel MYO3A missense mutation associated with a dominant form of late onset hearing loss. Sci Rep 8, 8706, doi:10.1038/s41598-018-26818-2 (2018).

14 Li, P. et al. Knock-In Mice with Myo3a Y137C Mutation Displayed Progressive Hearing Loss and Hair Cell Degeneration in the Inner Ear. Neural Plast 2018, 4372913, doi:10.1155/2018/4372913 (2018).

15 Doll, J. et al. A novel missense variant in MYO3A is associated with autosomal dominant high-frequency hearing loss in a German family. Mol Genet Genomic Med, e1343, doi:10.1002/mgg3.1343 (2020).

16 Liu, X. Z. et al. Autosomal dominant non-syndromic deafness caused by a mutation in the myosin VIIA gene. Nat Genet 17, 268–269, doi:10.1038/ng1197-268 (1997).

17 Nal, N. et al. Mutational spectrum of MYO15A: the large N-terminal extension of myosin XVA is required for hearing. Hum Mutat 28, 1014–1019, doi:10.1002/humu.20556 (2007).

18 Schneider, M. E. et al. A new compartment at stereocilia tips defined by spatial and temporal patterns of myosin IIIa expression. J Neurosci 26, 10243–10252, doi:10.1523/JNEUROSCI.2812-06.2006 (2006).

19 Li, S. et al. Myosin-VIIa is expressed in multiple isoforms and essential for tensioning the hair cell mechanotransduction complex. Nat Commun 11, 2066, doi:10.1038/s41467-020-15936-z (2020).

20 Grati, M. & Kachar, B. Myosin VIIa and sans localization at stereocilia upper tip-link density implicates these Usher syndrome proteins in mechanotransduction. Proc Natl Acad Sci U S A 108, 11476–11481, doi:10.1073/pnas.1104161108 (2011).

21 Rzadzinska, A. K., Schneider, M. E., Davies, C., Riordan, G. P. & Kachar, B. An actin molecular treadmill and myosins maintain stereocilia functional architecture and self-renewal. J Cell Biol 164, 887–897, doi:10.1083/jcb.200310055 (2004).

22 Masters, T. A., Kendrick-Jones, J. & Buss, F. Myosins: Domain Organisation, Motor Properties, Physiological Roles and Cellular Functions. Handb Exp Pharmacol 235, 77–122, doi:10.1007/164_2016_29 (2017).

23 Kazmierczak, P. et al. Cadherin 23 and protocadherin 15 interact to form tip-link filaments in sensory hair cells. Nature 449, 87–91, doi:10.1038/nature06091 (2007).

24 Assad, J. A., Shepherd, G. M. & Corey, D. P. Tip-link integrity and mechanical transduction in vertebrate hair cells. Neuron 7, 985–994, doi:10.1016/0896-6273(91)90343-x (1991).

25 Ahmed, Z. M. et al. The tip-link antigen, a protein associated with the transduction complex of sensory hair cells, is protocadherin-15. J Neurosci 26, 7022–7034, doi:10.1523/JNEUROSCI.1163-06.2006 (2006).

26 Webb, S. W. et al. Regulation of PCDH15 function in mechanosensory hair cells by alternative splicing of the cytoplasmic domain. Development 138, 1607–1617, doi:10.1242/dev.060061 (2011).

27 Michel, V. et al. Interaction of protocadherin-15 with the scaffold protein whirlin supports its anchoring of hair-bundle lateral links in cochlear hair cells. Sci Rep 10, 16430, doi:10.1038/s41598-020-73158-1 (2020).

28 Pepermans, E. et al. The CD2 isoform of protocadherin-15 is an essential component of the tip-link complex in mature auditory hair cells. EMBO Mol Med 6, 984–992, doi:10.15252/emmm.201403976 (2014).

29 Ebrahim, S. et al. Stereocilia-staircase spacing is influenced by myosin III motors and their cargos espin-1 and espin-like. Nat Commun 7, 10833, doi:10.1038/ncomms10833 (2016).

30 Cirilo, J. A., Jr., Gunther, L. K. & Yengo, C. M. Functional Role of Class III Myosins in Hair Cells. Front Cell Dev Biol 9, 643856, doi:10.3389/fcell.2021.643856 (2021).

31 Belyantseva, I. A., Boger, E. T. & Friedman, T. B. Myosin XVa localizes to the tips of inner ear sensory cell stereocilia and is essential for staircase formation of the hair bundle. Proc Natl Acad Sci U S A 100, 13958–13963, doi:10.1073/pnas.2334417100 (2003).

32 Senften, M. et al. Physical and functional interaction between protocadherin 15 and myosin VIIa in mechanosensory hair cells. J Neurosci 26, 2060–2071, doi:10.1523/JNEUROSCI.4251-05.2006 (2006).

33 Quintero, O. A. et al. Myosin 3A kinase activity is regulated by phosphorylation of the kinase domain activation loop. J Biol Chem 288, 37126–37137, doi:10.1074/jbc.M113.511014 (2013).

34 Dose, A. C. et al. The kinase domain alters the kinetic properties of the myosin IIIA motor. Biochemistry 47, 2485–2496, doi:10.1021/bi7021574 (2008).

35 Umeki, N. et al. The tail binds to the head-neck domain, inhibiting ATPase activity of myosin VIIA. Proc Natl Acad Sci U S A 106, 8483–8488, doi:10.1073/pnas.0812930106 (2009).

36 Bohil, A. B., Robertson, B. W. & Cheney, R. E. Myosin-X is a molecular motor that functions in filopodia formation. Proc Natl Acad Sci U S A 103, 12411–12416, doi:10.1073/pnas.0602443103 (2006).

37 Corey, E. J., Matsuda, S. P. & Bartel, B. Molecular cloning, characterization, and overexpression of ERG7, the Saccharomyces cerevisiae gene encoding lanosterol synthase. Proc Natl Acad Sci U S A 91, 2211–2215, doi:10.1073/pnas.91.6.2211 (1994).

38 Lin, C. H., Espreafico, E. M., Mooseker, M. S. & Forscher, P. Myosin drives retrograde F-actin flow in neuronal growth cones. Biol Bull 192, 183–185, doi:10.2307/1542600 (1997).

39 Mecklenburg, K. L. et al. Invertebrate and vertebrate class III myosins interact with MORN repeat-containing adaptor proteins. PLoS One 10, e0122502, doi:10.1371/journal.pone.0122502 (2015).

40 Senften, M. et al. Physical and Functional Interaction between Protocadherin 15 and Myosin VIIa in Mechanosensory Hair Cells. The Journal of Neuroscience 26, 2060–2071, doi:10.1523/jneurosci.4251-05.2006 (2006).

41 Tokuo, H., Mabuchi, K. & Ikebe, M. The motor activity of myosin-X promotes actin fiber convergence at the cell periphery to initiate filopodia formation. J Cell Biol 179, 229–238, doi:10.1083/jcb.200703178 (2007).

42 Sakai, T., Umeki, N., Ikebe, R. & Ikebe, M. Cargo binding activates myosin VIIA motor function in cells. Proc Natl Acad Sci U S A 108, 7028–7033, doi:10.1073/pnas.1009188108 (2011).

43 Wang, A. et al. Association of unconventional myosin MYO15 mutations with human nonsyndromic deafness DFNB3. Science 280, 1447–1451, doi:10.1126/science.280.5368.1447 (1998).

44 Mburu, P. et al. Defects in whirlin, a PDZ domain molecule involved in stereocilia elongation, cause deafness in the whirler mouse and families with DFNB31. Nat Genet 34, 421–428, doi:10.1038/ng1208 (2003).

45 Hadi, S., Alexander, A. J., Velez-Ortega, A. C. & Frolenkov, G. I. Myosin-XVa Controls Both Staircase Architecture and Diameter Gradation of Stereocilia Rows in the Auditory Hair Cell Bundles. J Assoc Res Otolaryngol 21, 121–135, doi:10.1007/s10162-020-00745-4 (2020).

46 Lu, Q., Li, J. & Zhang, M. Cargo recognition and cargo-mediated regulation of unconventional myosins. Acc Chem Res 47, 3061–3070, doi:10.1021/ar500216z (2014).

47 Salles, F. T. et al. Myosin IIIa boosts elongation of stereocilia by transporting espin 1 to the plus ends of actin filaments. Nat Cell Biol 11, 443–450, doi:10.1038/ncb1851 (2009).

48 Liu, H. et al. Myosin III-mediated cross-linking and stimulation of actin bundling activity of Espin. Elife 5, doi:10.7554/eLife.12856 (2016).

49 Rzadzinska, A., Schneider, M., Noben-Trauth, K., Bartles, J. R. & Kachar, B. Balanced levels of Espin are critical for stereociliary growth and length maintenance. Cell Motil Cytoskeleton 62, 157–165, doi:10.1002/cm.20094 (2005).

50 Zhao, B. et al. TMIE is an essential component of the mechanotransduction machinery of cochlear hair cells. Neuron 84, 954–967, doi:10.1016/j.neuron.2014.10.041 (2014).

51 Waguespack, J., Salles, F. T., Kachar, B. & Ricci, A. J. Stepwise morphological and functional maturation of mechanotransduction in rat outer hair cells. J Neurosci 27, 13890–13902, doi:10.1523/JNEUROSCI.2159-07.2007 (2007).

52 Porter, J. A., Yu, M., Doberstein, S. K., Pollard, T. D. & Montell, C. Dependence of calmodulin localization in the retina on the NINAC unconventional myosin. Science 262, 1038–1042, doi:10.1126/science.8235618 (1993).

53 Wolenski, J. S. Regulation of calmodulin-binding myosins. Trends Cell Biol 5, 310–316, doi:10.1016/s0962-8924(00)89053-4 (1995).

54 Dalal, J. S. et al. Mouse class III myosins: kinase activity and phosphorylation sites. J Neurochem 119, 772–784, doi:10.1111/j.1471-4159.2011.07468.x (2011).

55 Goldring, A. C., Beurg, M. & Fettiplace, R. The contribution of TMC1 to adaptation of mechanoelectrical transduction channels in cochlear outer hair cells. J Physiol 597, 5949–5961, doi:10.1113/JP278799 (2019).

56 McGrath, J. et al. Actin at stereocilia tips is regulated by mechanotransduction and ADF/cofilin. Curr Biol 31, 1141–1153.e1147, doi:10.1016/j.cub.2020.12.006 (2021).

57 Xiong, W. et al. TMHS is an integral component of the mechanotransduction machinery of cochlear hair cells. Cell 151, 1283–1295, doi:10.1016/j.cell.2012.10.041 (2012).

58 Maeda, R. et al. Tip-link protein protocadherin 15 interacts with transmembrane channel-like proteins TMC1 and TMC2. Proc Natl Acad Sci U S A 111, 12907–12912, doi:10.1073/pnas.1402152111 (2014).

59 Friedman, T. B., Sellers, J. R. & Avraham, K. B. Unconventional myosins and the genetics of hearing loss. Am J Med Genet 89, 147–157, doi:10.1002/(sici)1096-8628(19990924)89:3<147::aid-ajmg5>3.0.co;2-6 (1999).

60 Gibson, F. et al. A type VII myosin encoded by the mouse deafness gene shaker-1. Nature 374, 62–64, doi:10.1038/374062a0 (1995).

61 Walsh, T. et al. From flies’ eyes to our ears: mutations in a human class III myosin cause progressive nonsyndromic hearing loss DFNB30. Proc Natl Acad Sci U S A 99, 7518–7523, doi:10.1073/pnas.102091699 (2002).

62 Alagramam, K. N. et al. The mouse Ames waltzer hearing-loss mutant is caused by mutation of Pcdh15, a novel protocadherin gene. Nat Genet 27, 99–102, doi:10.1038/83837 (2001).

63 Naz, S. et al. Mutations of ESPN cause autosomal recessive deafness and vestibular dysfunction. J Med Genet 41, 591–595, doi:10.1136/jmg.2004.018523 (2004).

64 Schneider, C. A., Rasband, W. S. & Eliceiri, K. W. NIH Image to ImageJ: 25 years of image analysis. Nat Methods 9, 671–675, doi:10.1038/nmeth.2089 (2012).

65 Cording, J. et al. In tight junctions, claudins regulate the interactions between occludin, tricellulin and marvelD3, which, inversely, modulate claudin oligomerization. J. Cell Sci. 126, 554–564, doi:10.1242/jcs.114306 (2013).

